# Actin dependent membrane polarization reveals the mechanical nature of the neuroblast polarity cycle

**DOI:** 10.1101/2021.01.15.426888

**Authors:** Bryce LaFoya, Kenneth E. Prehoda

## Abstract

The Par complex directs fate determinant segregation from the apical membrane of asymmetrically dividing *Drosophila* neuroblasts. While the physical interactions that recruit the Par complex have been extensively studied, little is known about how the membrane itself behaves during polarization. We examined the membrane dynamics of neuroblasts and surrounding cells using a combination of super-resolution and time lapse imaging, revealing cellular-scale movements of diverse membrane features during asymmetric division cycles. Membrane domains that are distributed across the neuroblast membrane in interphase become polarized in early mitosis, where they mediate formation of cortical patches of the Par protein aPKC. Membrane and protein polarity cycles are precisely synchronized and are generated by extensive actin dependent forces that deform the surrounding tissue. In addition to suggesting a role for the membrane in asymmetric division, our results reveal the mechanical nature of the neuroblast polarity cycle.

## Introduction

The Par polarity complex mediates functions such as directional transport and fate determinant segregation in diverse animal cells (Lang and Munro, 2017; Riga et al., 2020; Venkei and Yamashita, 2018). A key step in Par complex function is formation of the Par domain, a continuous region along the cell membrane containing the Par complex proteins Par-6 and atypical Protein Kinase C (aPKC). Formation of the Par domain is a highly dynamic process involving recruitment from the cytoplasm and movement along the membrane (Gubieda et al., 2020; Illukkumbura et al., 2019; Sunchu and Cabernard, 2020). In asymmetrically dividing *Drosophila* neuroblasts, for example, initially cytoplasmic Par-6 and aPKC accumulate in the membrane’s apical hemisphere early in mitosis (Wodarz et al., 2000; Petronczki and Knoblich, 2001; Rolls et al., 2003; Homem and Knoblich, 2012). Membrane recruitment is followed by coalescence of the proteins into a cortical region concentrated around the apical pole (Oon and Prehoda, 2019). Extensive effort has been directed towards deciphering the physical interactions that recruit Par-6 and aPKC from the cytoplasm to the membrane. Here we examine the dynamics of the neuroblast membrane, both with the goal of understanding its potential role in the polarity cycle, but also to determine whether membrane dynamics might provide insight into the mechanism of formation of the Par domain.

While current models for neuroblast polarity focus on the protein-protein interactions that recruit the Par complex to the membrane (Knoblich, 2010; Lang and Munro, 2017; Venkei and Yamashita, 2018), accumulating evidence indicates that additional mechanisms may contribute to polarity. Neuroblast polarization is a stepwise process in which membrane targeting of the Par protein aPKC to the apical hemisphere is followed by coordinated movements towards the apical pole (Oon and Prehoda, 2019). Coalescence of aPKC requires the actin cytoskeleton (Oon and Prehoda, 2019), as does maintenance of the polarized state (Hannaford et al., 2018; Oon and Prehoda, 2019). However, the role of the actin cytoskeleton in the process remains unclear, as myosin II has been reported to be dispensable for apical polarity (Barros et al., 2003), and instead directly participates in polarization of the basal cortex (Barros et al., 2003; Hannaford et al., 2018). The potential lack of a role for myosin II in apical neuroblast polarity suggests that the actin cytoskeleton could play a passive role in contrast to the *C. elegans* zygote, for example, where actomyosin-generated cortical flows are essential for segregation of the Par complex along the membrane (Munro et al., 2004; Wang et al., 2017). Thus, while the actin cytoskeleton is required for neuroblast polarity, the extent to which force generation plays a role in the polarity cycle has been unclear.

Although the membrane plays a central role in Par-mediated polarity by establishing the scaffold for formation of the Par domain, its precise role in polarity remains unclear. In polarized cells such as epithelia, localized accumulations of specific phospholipids are thought to be important for formation of the Par domain, although the existence and function of these microdomains is controversial (van IJzendoorn et al., 2020; Riga et al., 2020; Stone et al., 2017). In the *C. elegans* zygote, a clear membrane asymmetry emerges during formation of the Par domain with filopodia-like structures preferentially forming in the anterior domain (Scholze et al., 2018; Hirani et al., 2019). What little is known about the neuroblast membrane suggest that it is relatively featureless as several phosphoinositide sensors have been reported to localize uniformly to the membrane (Doyle et al., 2017; Koe et al., 2018; Loyer and Januschke, 2018). Here we investigate whether the neuroblast membrane contains any heterogeneities and if so, whether these features might provide insight into the polarity cycle.

## Results

### The neuroblast membrane is heterogeneous and forms extensive contacts with progeny cells

We examined the neuroblast membrane using three markers, each of which interacts with the membrane via a distinct mechanism: a farnesyl-modified peptide that integrates into the bilayer (FP-farnesyl) (Zhou et al., 2015), the pleckstrin homology domain from PLCδ that interacts with the headgroup of the phosphoinositide PI(4,5)P2 (FP-PH) (Verstreken et al., 2009), and the integral membrane protein CD8 that traverses the bilayer (FP-CD8) (Lee and Luo, 1999). We expressed fluorescent protein fusions of the markers specifically in the neuroblast lineage using the UAS promoter and worniu-GAL4 driver, and imaged actively dividing central nervous system neuroblasts from third instar larval brain explants (Figure 1A,B). To image neuroblast membranes at super-resolution, we used spinning-disk confocal microscopy with optical photon reassignment (Azuma and Kei, 2015). Each of the membrane markers outlined the plasma membrane of neuroblasts and their smaller progeny cells, with little background from other cells (Figure 1C and Video S1). For FP-farnesyl and FP-PH, the plasma membrane was marked nearly exclusively (small vesicle-like signals were occasionally seen with each) whereas FP-CD8 also marked some internal membranes.

**Figure 1.**
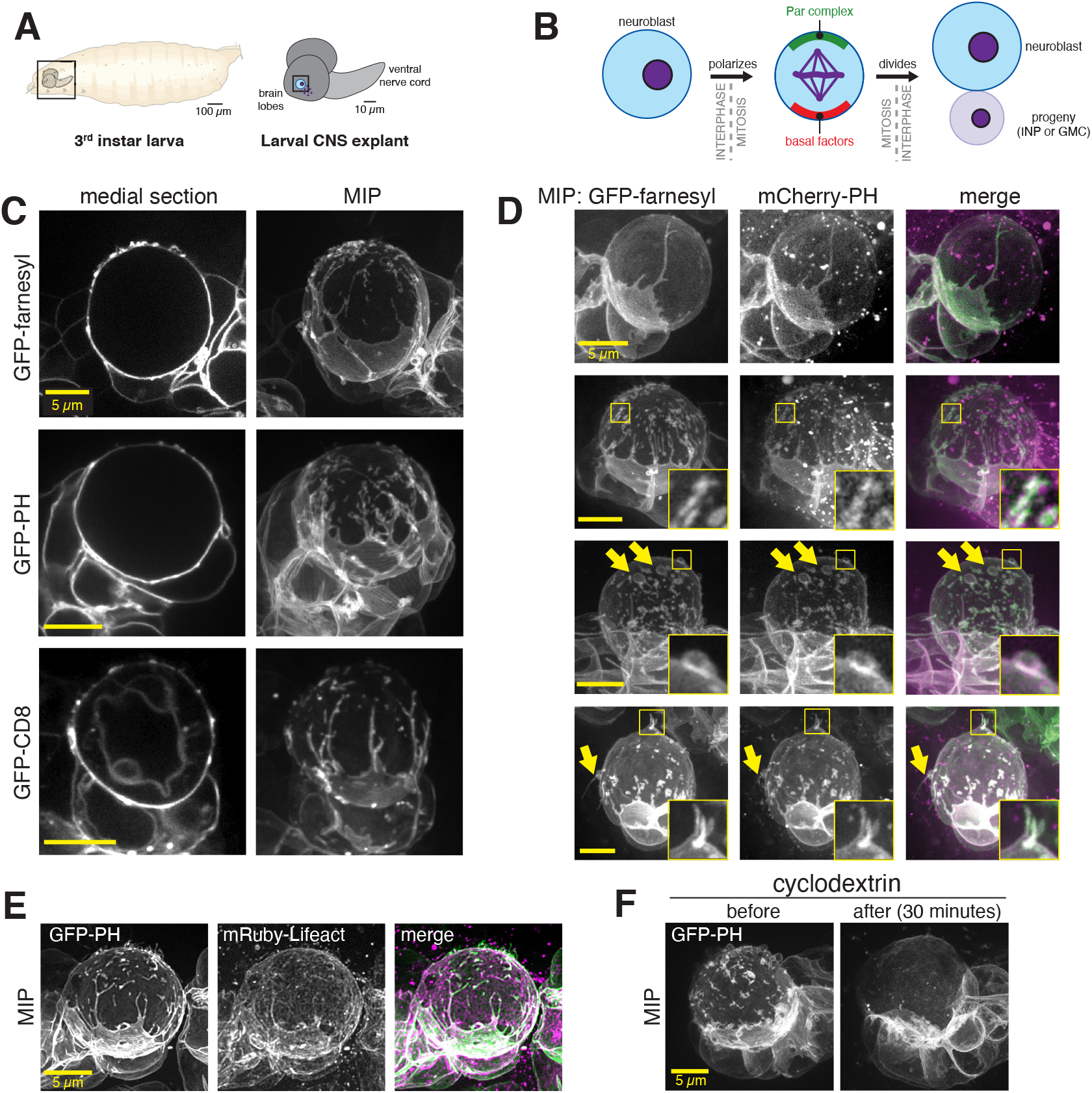
The neuroblast membrane is heterogenous. (**A**) *Drosophila* larval central nervous system explants used in this study. (**B**) Larval brain neuroblast polarity cycle. Division results in self-renewal of the apical daughter and production of an Intermediate Neuronal Precursor (INP) or Ganglion Mother Cell (GMC). (**C**) Super resolution images of neuroblasts and their progeny from *Drosophila* third instar larvae. Neuroblasts expressed the indicated membrane marker under the control of the UAS promoter and driven by worniu-GAL4. A single medial optical section is shown along with a maximum intensity projection made from optical sections through the neuroblast’s front hemisphere (MIP). Rotations of each cell’s full volume MIP are shown in Video S1. (**D**) Direct comparison of neuroblast features using simultaneous imaging of multiple membrane markers. Images are maximum intensity projections as in (C). Selected features are highlighted with arrows and boxes with magnified inset. Rotations of each cell’s full volume MIP are shown in Video S2.(**E**) Colocalization of neuroblast membrane features (GFP-PH) and actin (mRuby-Lifeact). (**F**) Effect of cyclodextrin treatment on neuroblast membrane heterogeneity. Maximum intensity projections constructed from optical sections taken from the same neuroblast before and 30 minutes following treatment with cyclodextrin are shown.

Individual optical sections and maximum intensity projections (MIPs) of the full cell volume revealed a rich landscape of neuroblast membrane features, including blebs and filopodial-like extensions, smaller domains, and extensive contacts with progeny cells (Figure 1C,D and Videos S1 and S2). Extensions were typically greater than 0.5 μm in length whereas membrane domains were smaller areas of marker enrichment that were closer to the cell body but also protruded to some extent. The membrane markers also revealed that progeny cells form extensive contacts with their associated neuroblast. These contacts were large in surface area and also typically included filopodial extensions that emanated from the progeny cell towards the opposite side of the neuroblast. In many instances, the progeny filopodia encircled the neuroblast to an extent that the progeny appeared to engulf part of the neuroblast. We observed the same membrane features when simultaneously imaging distinct membrane markers (Figure 1D), indicating that the markers report on the same set of features. Furthermore, membrane features were associated with cortical F-actin, as detected by the Lifeact sensor (Figure 1E).

The smaller membrane domains dissipated when methyl-β-cyclodextrin was included in the surrounding media (Figure 1F; n = 25 neuroblasts). Cyclodextrin sensitivity and CD8 enrichment are characteristics of membrane microdomains – localized areas where specific phospholipids and proteins are concentrated (Barman and Nayak, 2007; Pang et al., 2007), although the increased membrane marker signal may be due to increased amounts of plasma membrane, consistent with their slight protrusion above the cell surface (Figure 1A-E) and similar observations in the worm zygote (Hirani et al., 2019).

### The neuroblast membrane undergoes an actin cytoskeleton-dependent polarity cycle

To examine neuroblast membrane dynamics, we acquired time series of optical sections across the cell volume approximately once per five minutes, a frequency which did not cause significant photobleaching using the optical reassignment imaging method. Even at this temporal resolution, maximum intensity projections of sections through the cell’s full volume revealed changes in the membrane over the course of the cell cycle (Figure 2A). Membrane dynamics were not constant, with relatively little change until sometime in mitosis when large movements appeared to begin before cleavage furrow ingression and end shortly after the completion of division.

**Figure 2.**
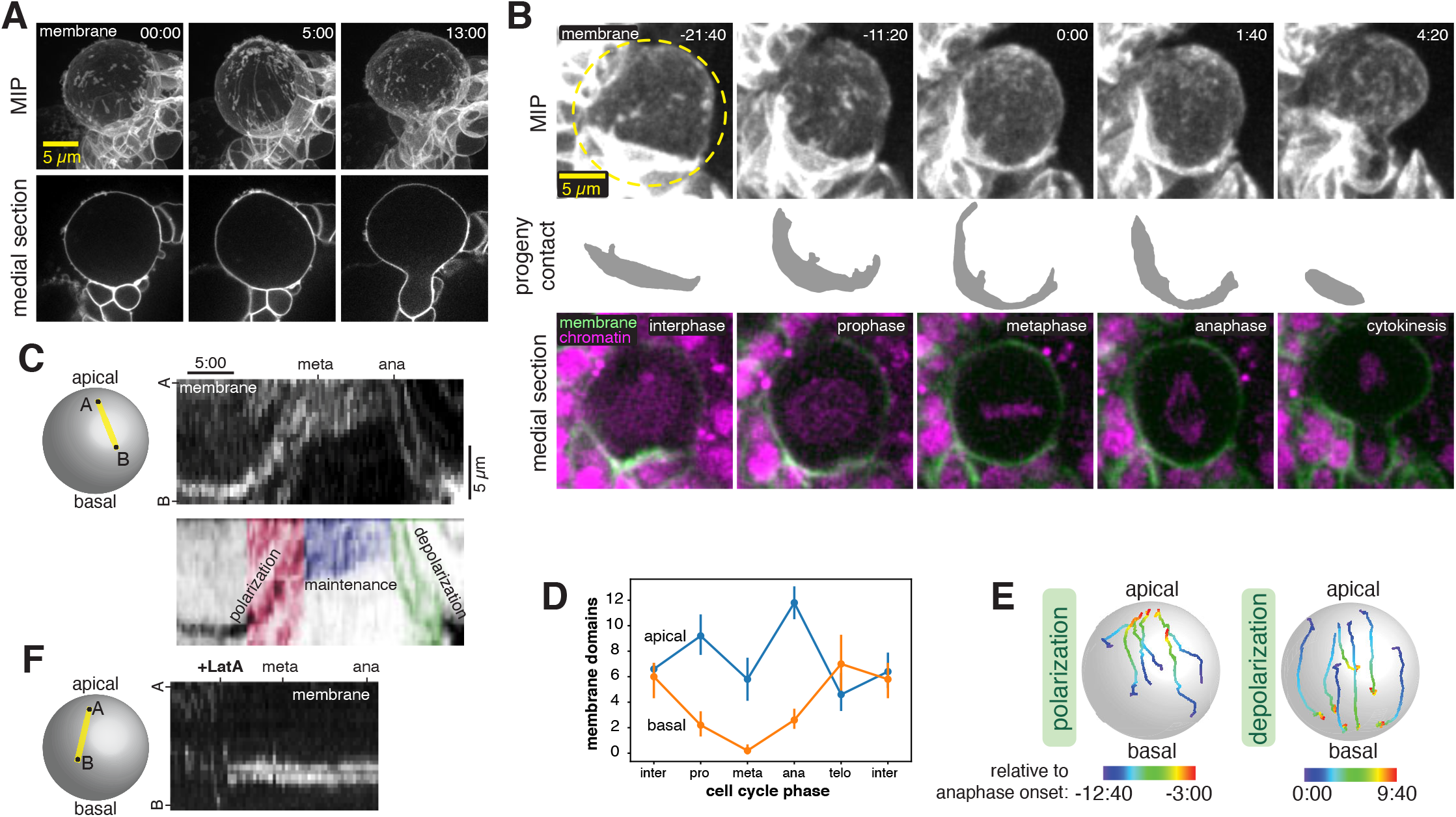
The neuroblast membrane undergoes a polarity cycle. (**A**) Membrane dynamics of a dividing neuroblast and progeny cells observed with super resolution imaging of GFP-farnesyl (”membrane”). A single medial optical section is shown along with a maximum intensity projection (MIP) of optical sections through the front hemisphere. Time in minutes and seconds relative to when the first frame was collected is shown in each frame. (**B**) Neuroblast membrane dynamics through a full division cycle. Selected frames from Video S3 are shown in maximum intensity projection of the front hemisphere GFP-PH signal (”membrane”), with a manual segmentation of the basal daughter cell membrane that contacts the neuroblast (progeny contact), and a medial section with both GFP-PH and RFP-H2A (”chromatin”) signals. Time in minutes and seconds relative to anaphase onset is indicated in the upper right corner of each frame. (**C**) Kymograph along the indicated axis following the progression of a GFP-PH membrane domain over the course of mitosis (from Video S3). A legend showing the three phases of dynamics is shown below the kymograph. (**D**) Membrane domain count in the apical and basal hemispheres at different cell cycle phases (n = 5 different neuroblasts). Note that the domain count at metaphase decreases because the apical domains coalesce. (**E**) Particle tracking of individual membrane domains during both the polarization and depolarization phases from Video S3. The color of each track is coded by the time relative to anaphase onset. (**F**) Kymograph along the indicated axis following the progression of a GFP-PH membrane domain over the course of mitosis in a LatA treated neuroblast (from Video S4A).

To follow the neuroblast membrane at higher temporal resolution, we used standard resolution spinning disk confocal microscopy, imaging the full cell volume every 20 seconds. Additionally, we simultaneously imaged the chromosomal marker RFP-His2A to more precisely identify the cell cycle stages at which transitions in membrane dynamics occur. From the resulting movies we identified four distinct temporal phases of neuroblast membrane movements (Figure 2B,C and Video S3). In interphase, membrane features exhibited only small movements that were uncoordinated and lacked any clear directionality. The relative calm of the interphase membrane gave way to a period of highly coordinated dynamics in early prophase in which features across the cell membrane moved rapidly towards the apical pole. We detected apical movement not only in features such as localized marker enrichments, but in progeny cell contacts that are typically near the basal pole, indicating that substantial force accompanies these movements (Figure 2B and Video S3). Continued apical movement during prophase ultimately led to accumulation of membrane features in the apical hemisphere of the cell (i.e. polarization) until this phase ends near metaphase (Figure 2B-E).

The polarized membrane state was maintained for a short period until anaphase began and membrane movements reversed direction, leading to their dispersal across the membrane surface (i.e. depolarization) by the completion of cytokinesis. The basally-directed movements that occur in anaphase also caused the progeny cell contacts at the basal region of the neuroblast membrane to relax to their pre-polarization state (Figure 2B and Video S3). Thus, there are four phases of membrane dynamics during neuroblast asymmetric cell division: 1. a relatively static interphase with features evenly distributed across the cell surface, 2. apically-directed movement during prophase that generates an 3. apically polarized state that is maintained through metaphase, and 4. a depolarization phase during anaphase. The process that drives membrane dynamics operates at the cellular scale, altering both the basal and apical membranes, and the force generated during the process significantly deforms the surrounding tissue.

To gain insight into the underlying process that drives membrane movements, we examined membrane dynamics in neuroblasts treated with the actin depolymerizing drug Latrunculin A (LatA). As shown in Figure 2F and Video S4, membrane domains halt apically-directed movements during prophase immediately following LatA exposure and become nearly completely static (n = 15 neuroblasts). However, while the dynamics of membrane domains were dependent on the actin cytoskeleton, the domains themselves were unaffected by LatA treatment. Simultaneous imaging of GFP-PH and mRuby-Lifeact confirmed that membrane movements ceased when cortical actin dissipated (Video S4). Thus, an intact actin cytoskeleton is required for the complex dynamics of membrane domains during asymmetric cell division, but the domains persist in the absence of F-actin.

### The Par protein aPKC localizes to membrane domains and follows their dynamics

The phases of neuroblast membrane dynamics we observed, rapid polarization and then depolarization, resemble those of Par polarity proteins such as Bazooka (Baz; aka Par-3) and aPKC (Oon and Prehoda, 2019), with the key difference that membrane domains are present throughout the cell cycle whereas aPKC is cytoplasmic in interphase and does not begin accumulating on the membrane until early prophase (Hannaford et al., 2018; Oon and Prehoda, 2019). We first examined the localization of the aPKC and Baz using super-resolution imaging and found that while aPKC significantly overlapped with the membrane domains, Baz was predominantly localized away from domains (Figure 3A,B). We therefore examined whether membrane and aPKC dynamics are correlated by imaging larval brain neuroblasts expressing GFP-aPKC from its endogenous promoter and *worniu-GAL4* driven mCherry-PH. We observed accumulation of aPKC early in mitosis, with localized enrichments (i.e. patches) forming at apically localized membrane domains and aPKC that was more diffusely distributed elsewhere in the apical hemisphere (Figure 3 and Video S5). Both pools of cortical aPKC, membrane domain-localized patches and diffuse, moved towards the apical pole simultaneously, with protein and membrane movement beginning simultaneously (i.e. within one frame; n = 8 neuroblasts). The membrane domains often appeared to encircle the aPKC (Figure 3A,C and Video S5) and membrane domains that were initially in the basal hemisphere did not recruit aPKC until they passed the equator into the apical hemisphere (Figure 3E and Video S6; n = 3).

**Figure 3.**
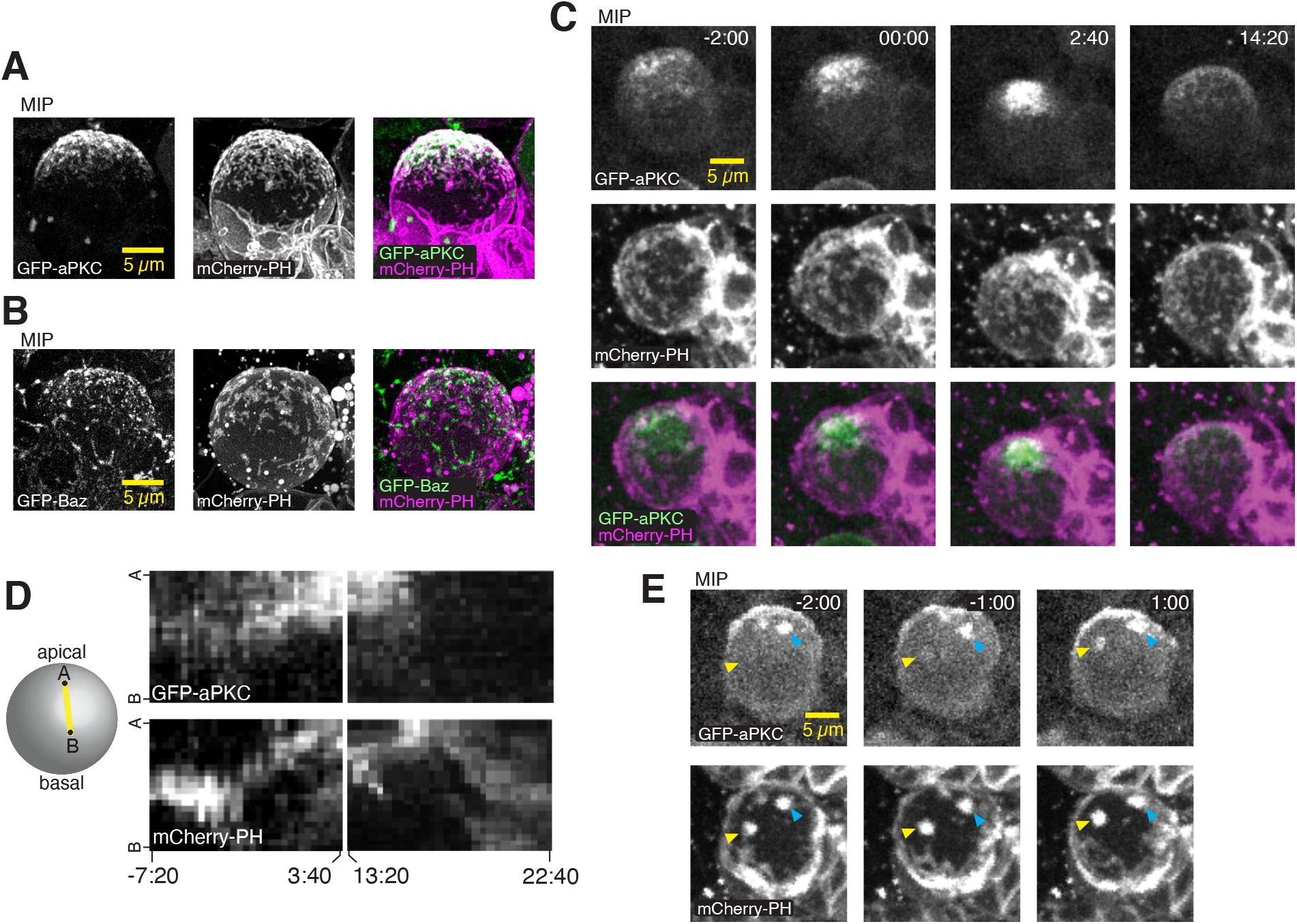
Neuroblast membrane and polarity protein dynamics are coupled. (**A**) Maximum intensity projection (MIP) constructed from super resolution optical sections of the polarity proteins aPKC and the mCherry-PH membrane marker. (**B**) MIP of Bazooka (Baz; aka Par-3) and mCherry-PH as in (A). (**C**) Simultaneous imaging of membrane and aPKC dynamics during neuroblast asymmetric division. Selected frames from Video S5 of a neuroblast expressing GFP-aPKC from its endogenous promoter and mCherry-PH via worniu-GAL4. Time relative to nuclear envelope breakdown (NEB) is shown. As described previously, aPKC accumulates on the apical membrane beginning in early prophase followed by a coalescence phase shortly before NEB. Membrane features are present throughout the cell cycle, and their movements occur simultaneously with aPKC. (**D**) Kymograph along the apical-basal axis of Video S5 showing the correlated dynamics of aPKC and the neuroblast membrane during polarization and depolarization. (**E**) Selected frames from Video S6 showing an example of aPKC recruitment to a membrane patch as it moves from the basal to the apical hemisphere. Yellow arrowheads mark the membrane patch and the corresponding aPKC signal. Cyan arrowheads mark a membrane patch and corresponding aPKC signal that starts and ends in the apical hemisphere.

Upon completion of the polarization process, membrane domains and aPKC remained tightly localized around the apical pole for a short period before disassembling via basally directed movements. Disassembly of the polarized membrane domains and aPKC began simultaneously (within one frame; n = 8 neuroblasts), as both moved rapidly towards the emerging cleavage furrow. Following disassembly, aPKC signal rapidly dissipated from the membrane, whereas the depolarized membrane domains persisted into the following interphase. Thus, aPKC patches form at apically localized membrane domains and both populations of cortical aPKC, patches and diffuse, precisely follow membrane dynamics with the same coordinated apically- and basally-directed movements during prophase and anaphase, respectively.

### Neuroblast membrane domains mediate aPKC cortical patch formation

The targeting of aPKC cortical patches to membrane domains and the correlated movement of membrane and polarity protein suggest that the two may be functionally related. We tested the effect of removing membrane domains on the cortical localization of aPKC by examining aPKC localization in neuroblasts that lacked membrane domains owing to cyclodextrin treatment. These neuroblasts failed to form patches of aPKC, with the remaining cortical aPKC signal diffusely distributed over the apical cortex (n = 12; Figure 4A and Video S7). The diffuse aPKC continued to polarize at the apical pole in the absence of membrane domains as it did in untreated neuroblasts (Figure 3 and Video S5) although the overall signal was significantly reduced in the cyclodextrin-treated neuroblasts (Figure 4B). We also tested whether aPKC is required for membrane domains to form and noticed no effect on membrane structure in neuroblasts expressing an RNAi directed against aPKC (Figure 4C and Video S8).

**Figure 4.**
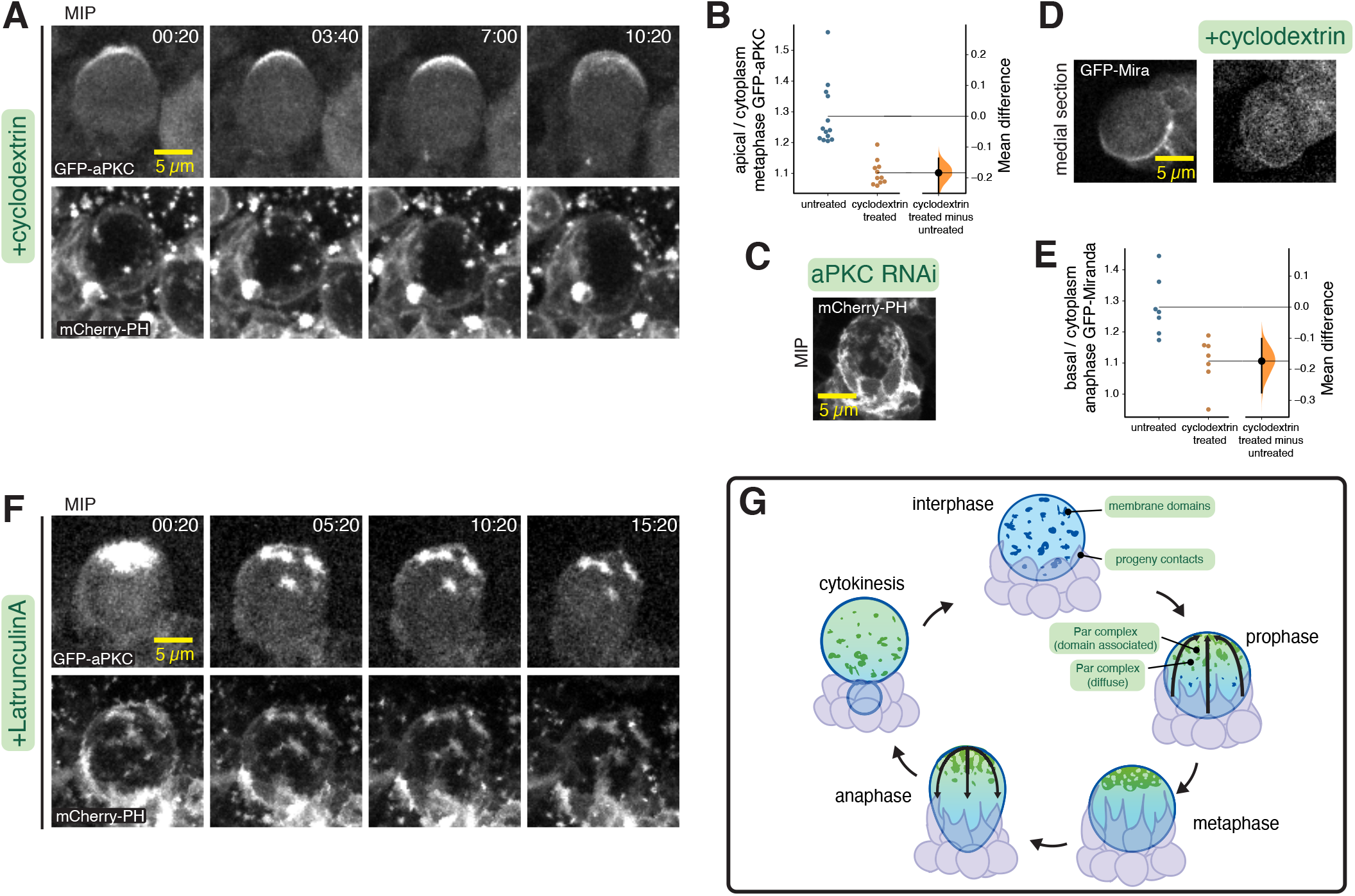
Membrane domains mediate aPKC polarity maintenance. (**A**) Membrane and aPKC dynamics in a neuroblast in which membrane domains were removed via cyclodextrin treatment. Selected frames subsequent to polarization are shown from Video S7. Time is shown in minutes and seconds relative to NEB. (**B**) Gardner-Altman estimation plot of the effect of cyclodextrin on the cortical recruitment of aPKC. Apical cortical to cytoplasmic signal intensity ratios of GFP-aPKC are shown for individual metaphase neuroblasts treated with cyclodextrin. Statistics: bootstrap 95% confidence interval. (**C**) Neuroblast expressing mCherry-PH and aPKC RNAi. Selected frame from Video S8. (**D**) Comparison of GFP-Miranda in untreated and cyclodextrin-treated neuroblasts (anaphase) shown in medial sections. MIPs for the same neuroblasts are shown in Video S9. (**E**) Gardner-Altman estimation plot of the effect of cyclodextrin on the cortical recruitment of Miranda. Basal cortical to cytoplasmic signal intensity ratios of GFP-Miranda are shown for individual anaphase neuroblasts treated with cyclodextrin. Statistics: bootstrap 95% confidence interval. (**F**) Membrane and aPKC maintenance dynamics in a neuroblast in which the actin cytoskeleton was disrupted by treatment with LatA. Selected frames following LatA addition are shown from S10. (**G**) Model for neuroblast membrane dynamics. Membrane features are dispersed over the entire cell surface during interphase and progeny cell contacts are typically restricted to the basal pole. Apically-directed movements begin in prophase, when aPKC is recruited to the apical hemisphere of the membrane (enriched at membrane domains and diffuse elsewhere), and movements lead to metaphase polarization and deformation of progeny contacts. Basally-directed movements begin in anaphase and depolarize membrane features and return progeny cell contacts to the basal hemisphere.

To determine if the loss of cortical patches influenced aPKC’s ability to regulate polarity, we examined the localization of the basal factor Miranda in cyclodextrin-treated neuroblasts. Miranda is normally polarized to the basal cortex due to the activity of aPKC, leading to its segregation into the basal daughter cell during cytokinesis (Rolls et al., 2003). Although Miranda polarization was still detectable in cyclodextrin-treated neuroblasts, its basal localization at anaphase was significantly reduced compared to untreated neuroblasts (Figure 4D,E and Video S9). Taken together, the effects of cyclodextrin on aPKC and Miranda indicate that membrane domains are required for cortical aPKC patches, and that aPKC and Miranda polarity are significantly reduced without patches. Importantly, the diffusely cortical aPKC that remains in the absence of membrane domains is polarized similarly (as are the progeny cell contacts) suggesting that the underlying process that drives coalescence does not require the domains.

The actin cytoskeleton is required to maintain the diffusely localized pool of cortical aPKC in the apical hemisphere (Hannaford et al., 2018; Oon and Prehoda, 2019). We examined whether the membrane domain-localized cortical aPKC patches also require F-actin to retain aPKC. In LatA-treated neuroblasts expressing GFP-aPKC and mCherry-PH as a membrane marker, we observed rapid dissipation of cortical aPKC that was not colocalized with a membrane domain (signal below limit of detection in 24.9 ± 10 minutes, n = 21; Figure 4F and Video S10), as previously described (Oon and Prehoda, 2019). However, aPKC patches remained localized to membrane domains for very extended time periods in all neuroblasts examined (38.7 ± 6 minutes – many neuroblasts had signal remaining at the end of the imaging session, n = 21), indicating that their retention in the apical hemisphere does not require the actin cytoskeleton. Thus, the two pools of cortical aPKC are maintained in the apical hemisphere via an actin-dependent mechanism for diffusely localized aPKC, and a membrane structure-dependent one for cortical aPKC patches.

## Discussion

We examined the neuroblast membrane using super resolution imaging and found that it is very heterogenous, with features such as small cyclodextrin-sensitive domains, filopodial extensions, and extensive contacts with progeny cells (Figure 1 and Videos S1,S2). Membrane domains are initially dispersed across the cell surface in interphase, with progeny contacts typically near the basal pole. The membrane undergoes several phases of coordinated movements during mitosis (Figure 4G), first polarizing by enriching features in the apical hemisphere with coordinated, apically-directed movements that are dependent on the actin cytoskeleton. The forces that generate the polarized neuroblast membrane deform the surrounding tissue, indicating that they are significant in magnitude and scale. After a brief phase where the polarized state is maintained, basally-directed movements depolarize the membrane by redistributing the features across the cell surface. We found that these complex dynamics, including the timing of the transitions between phases and the characteristics of the movements, are precisely correlated with those of the Par polarity protein aPKC (Figure 3 and Video S5). Furthermore, membrane domains participate in the recruitment and retention of aPKC, specifically recruiting aPKC when they are in the apical hemisphere (Figure 3E and Video S6), and removing domains with cyclodextrin significantly reduces both apical and basal polarity (Figure 4B,E).

The behavior of the membrane indicates that the neuroblast polarity cycle is a mechanical process in which cellular-scale forces are generated. Par proteins were recently found to undergo apically-directed movements that lead to neuroblast polarization (Oon and Prehoda, 2019), but whether the underlying cellular process that drives these movements acts locally or at a larger scale has not been known. We observed movement of membrane features in the apical hemisphere but also near the basal pole, including the extension of filopodial-like arms from progeny cells that wrap around the neuroblast. The deformation of the surrounding tissue raises the possibility that the polarity cycle could be driven by contacting cells rather than cell autonomously by the neuroblast, as has been observed in some epithelial cells (Pohl et al., 2012; Roh-Johnson et al., 2012). However, we believe two features of the data strongly support a cell autonomous model: membrane domains distant from progeny contacts move with the same dynamics, and the polarity cycle is tightly coupled to the neuroblast’s cell cycle (Figure 2 and Video S3).

The membranes of both the *Drosophila* neuroblast (present work) and the *C. elegans* zygote (Scholze et al., 2018; Hirani et al., 2019) are polarized during asymmetric division. The zygote membrane is initially devoid of features, and while some movement occurs, it is primarily polarized by the preferential appearance of filopodial like structures in the anterior domain. In contrast, features are present on the neuroblast membrane throughout the cell cycle and polarization results from the coordinated movement of membrane domains towards the apical pole (Figure 2). In the zygote, pulsatile contractions of actomyosin generate cortical flows important for Par polarity. We have found that an intact actin cytoskeleton is necessary for neuroblast membrane polarity (Figure 2). However, myosin II has been reported to be directly involved in basal, not apical neuroblast polarity (Barros et al., 2003; Hannaford et al., 2018), and while myosin II has been extensively imaged in the neuroblast (Barros et al., 2003; Cabernard et al., 2010; Connell et al., 2011; Koe et al., 2018; Roth et al., 2015; Roubinet et al., 2017; Tsankova et al., 2017), cortical actomyosin dynamics during polarization have not been reported. Future work will be directed at understanding how the extensive forces that occur during the neuroblast polarity cycle are generated.

Our results indicate that the neuroblast membrane plays a role in polarity initiation and maintenance. We observed recruitment of aPKC to apical membrane domains and retention of aPKC at these domains even in the absence of the actin cytoskeleton (Figures 3 and 4). While aPKC is also recruited to apical sites outside of membrane domains, this “diffuse” aPKC requires the actin cytoskeleton and rapidly depolarizes in the presence of LatA. Furthermore, when membrane domains are ablated with cyclodextrin, polarity is significantly reduced (Figure 4B,E). Thus, we propose that the two pools of membrane-bound aPKC, diffuse and membrane-domain associated, work together to initiate and maintain apical neuroblast polarity (Figure 4G). Given that the membrane density appears to be higher at domain sites (Figure 1), the extended maintenance at domains could arise simply from an initially higher concentration of aPKC. Further work will be necessary to understand if domain associated aPKC is mechanistically distinct from its diffuse counterpart.

## RESOURCE AVAILABILITY

### Lead Contact

Further information and requests for resources and reagents should be directed to the Lead Contact, Kenneth Prehoda.

### Materials Availability

This study did not generate new unique reagents.

### Data and Code Availability

The raw data supporting the current study have not been deposited in a public repository because of their large file size but are available from the corresponding author on request.

## EXPERIMENTAL MODEL AND SUBJECT DETAILS

### Fly Strains

For live imaging of membrane dynamics in neuroblasts, a Worniu-Gal4 driver line was used to express fluorescent membrane-bound fusion proteins under UAS control. Three classes of membrane-bound fusion proteins were used. FP-farnesyl expresses the C-terminal region of human K-Ras tagged with GFP which becomes farnesylated and membrane-anchored in cells. FP-PH expresses the pleckstrin homology domain of human PLCδ tagged with GFP or mCherry, which binds to the plasma membrane lipid phosphoinositide PI(4,5)P2. FP-CD8 (a gift from the Chris Doe Lab) expresses a single-pass transmembrane protein tagged with GFP. RFP-His2A expresses RFP-tagged His2A under the control of native promoters. The BAC-encoded GFP-aPKC (Besson et al., 2015) was used to track aPKC localization and dynamics.

## METHOD DETAILS

### Live Imaging

Intact *Drosophila* central nervous systems were dissected from third instar larvae in a bath of Schneider’s Insect Media. These larval brain explants were then mounted dorsal side down on sterile poly-D-lysine coated 35mm glass bottom dishes (ibidi Cat#81156) containing modified minimal hemolymph-like solution (HL3.1). Explants were imaged on a Nikon Eclipse Ti-2 (60x H_2_O objective) equipped with a Yokogawa CSU-W1 SoRa spinning disk head and dual Photometrics Prime BSI sCMOS cameras. GFP tagged proteins were illuminated with 488nm laser light. RFP, mCherry, and mRuby tagged proteins were illuminated with 561nm laser light. For time-lapse imaging 41-61 optical sections with a step size of 0.5 *μ*m were acquired every 20 seconds. For super resolution imaging, individual neuroblasts were imaged with a step size of 0.3 *μ*m using SoRa optics which achieve super resolution through optical photon reassignment. To examine the role of F-actin in membrane and aPKC dynamics, explants were treated with 50 *μ*M latrunculin A (0.5% DMSO) during imaging. To examine the role of cholesterol in membrane and aPKC dynamics, explants were treated with 15 mM methyl-B-cyclodextrin (solubilized in HL3.1) during imaging. We observed loss of F-actin within minutes of LatA treatment but membrane domains persisted approximately 30 minutes following cyclodextrin treatment, presumably because of cyclodextrin’s larger mass and concomitant slower diffusion into the tissue.

### Image Processing and Analysis

Images were analyzed using ImageJ (FIJI package) and Imaris (Bitplane) software. Neuroblasts were identified through their location in the brain and large size, as well as the presence of tissue-specifically expressed transgenes. For rotating movies, maximum intensity projections (MIPs) were assembled from optical slices through the entirety of the neuroblast volume. Photobleaching during time-lapse imaging was corrected for using ImageJ bleach correction tool in histogram matching mode. For aPKC images, a guassian blur of 0.5 pixels was used to improve signal to noise. For imaging of RFP and mCherry tagged transgenes, ImageJ despeckle and/or smooth tools were applied. Kymographs were created from time-lapse movies using a line along the basal-apical axis using ImageJ multi kymograph tool. To quantify the number of membrane domains in the apical and basal hemispheres of the membrane, larval neuroblasts (n=5) expressing worniu-GAL4 driven GFP-PH were imaged using traditional spinning disk microscopy. Membrane domain were distinguished based on a cutoff of 3X signal intensity above background and an area of at least 0.3 square micrometers. Membrane domains were found to range in size from 0.3-5 square micrometers. To quantify the effect of cyclodextrin on aPKC and Miranda recruitment, movies from larval neuroblasts expressing either GFP-aPKC from its native promoter or worniu-GAL4 driven GFP-Miranda were analyzed. For GFP-aPKC, medial optical slices from post NEB, pre-anaphase cells captured using SoRa spinning disk microscopy were then used to trace the apical cortex and averaged across this region of interest in FIJI. A second region of interest within the cytoplasm was used to determine the apical:cytoplasmic ratio for untreated (n=13) vs cyclodextrin treated (n=11) neuroblasts. A similar protocol was used to assess miranda signal intensity along the basal cortex at anaphase onset however medial slices acquired using traditional spinning disk time lapse imaging were used to trace the basal cortex from which miranda signal intensity was measured and averaged across this region of interest in FIJI. A second region of interest within the cytoplasm was used to determine the basal:cytoplasmic ratio for untreated (n=7) vs cyclodextrin treated (n=7) neuroblasts. Particle tracking was performed using the ‘spots’ tracking utility in Imaris. Here, membrane domains were tracked in a neuroblast expressing worniu-GAL4 driven GFP-PH during both the polarization and depolarization phases of the polarity cycle. (edited)

## QUANTIFICATION AND STATISTICAL ANALYSIS

Not applicable

## KEY RESOURCES TABLE

| REAGENT or RESOURCE | SOURCE | IDENTIFIER |
| --- | --- | --- |
| Chemicals, Peptides, and Recombinant Proteins |  |  |
| Latrunculin A | Enzo Life Sciences | Cat#BML-T119-0100, CAS# 76343-93-6 |
| Methyl-β-cyclodextrin | Caymen Chemical | Cat#21633, CAS# 128446-36-6 |
| Experimental Models: Organisms/Strains |  |  |
| UAS-aPKC RNAi | Vienna *Drosophila* Resource Center | v105624 |
| FP-Farnesyl | Bloomington *Drosophila* Stock Center | BDSC Cat#80052, RRID:BDSC_80052 |
| FP-PH (GFP) | Bloomington *Drosophila* Stock Center | BDSC Cat#39693, RRID:BDSC_39693 |
| FP-CD8 | Bloomington *Drosophila* Stock Center | BDSC Cat#5130, RRID:BDSC_5130 |
| Worniu-Gal4 | Chris Q. Doe Lab |  |
| RFP-His2a (H2A) | Bloomington *Drosophila* Stock Center | BDSC Cat#23650, RRID:BDSC_23650 |
| GFP-aPKC | François Schweisguth Lab; Besson et al., 2015 |  |
| FP-PH (mCherry) | Bloomington *Drosophila* Stock Center | BDSC Cat#51658,<br>RRID:BDSC_51658 |
| GFP-Miranda | Bloomington *Drosophila* Stock Center | BDSC Cat#56555,<br>RRID:BDSC_56555 |
| mRuby-Lifeact | Bloomington *Drosophila* Stock Center | BDSC Cat#35545,<br>RRID:BDSC_35545 |
| Baz-GFP | Carnegie Protein Trap Library | PMID: 17194782 |
| Software and Algorithms |  |  |
| ImageJ (FIJI package) | National Institutes of Health |  |
| Imaris | Bitplane |  |

## Video legends

Video S1: Maximum intensity projections of neuroblasts and progeny cells expressing membrane markers.

Super resolution images of neuroblasts and their progeny from *Drosophila* third instar larvae. Neuroblasts expressed the indicated membrane marker under the control of the UAS promoter and driven by worniu-GAL4. Maximum intensity projections (MIP) made from optical sections through the neuroblast’s entire volume are shown. Rotations of each cell’s full volume MIP are shown in the following order: GFP-farnesyl, GFP-PH, GFP-CD8.

Video S2: The neuroblast membrane is a rich landscape of features.

Super resolution images of neuroblasts and their progeny from Drosophila third instar larvae. Neuroblasts expressed GFP-farnesyl under control of the UAS promoter and driven by worniu-GAL4. Rotating of maximum intensity projections (MIP) made from optical sections through the neuroblast’s entire volume are shown.

Video S3: Neuroblast membrane dynamics through a full division cycle.

Time lapse imaging of neuroblasts and their progeny from Drosophila third instar larvae through a complete division cycle. Neuroblasts expressed GFP-PH under control of the UAS promoter driven by worniu-GAL4, and RFP-H2A under control of a constitutive promoter. Left: maximum intensity projection of the front hemisphere GFP-PH signal (”membrane”). Right: medial section with both GFP-PH in green and RFP-H2A (“chromatin”) in magenta. Time relative to anaphase onset (mm:ss) is shown.

Video S4: Membrane dynamics in neuroblasts treated with the actin depolymerizing drug Latrunculin A.

Time lapse imaging of neuroblasts and their progeny from *Drosophila* third instar larvae through a complete division cycle in the presence of LatA. The first two movies are of neuroblasts which expressed GFP-PH under control of the UAS promoter driven by worniu-GAL4 and RFP-H2A under control of a constitutive promoter. Movie A begins with the neuroblast in an untreated state before LatA was added to the media. Caption denotes when LatA was added. Left: maximum intensity projection of the front hemisphere GFP-PH signal (”membrane”). Right: medial section with GFP-PH in green and RFP-H2A (”chromatin”) in magenta. Movie B shows the same genotype and imaging conditions as the first movie, but was treated DMSO, serving as the vehicle control. Movie C shows a neuroblast which expressed GFP-PH and mRuby-Lifeact both under control of the UAS promoter driven by worniu-GAL4. The movie begins with the neuroblast in a untreated state before LatA was added to the media. Caption denotes when LatA was added. Left: maximum intensity projection of the front hemisphere GFP-PH signal. Middle: maximum intensity projection of the front hemisphere mRuby-Lifeact signal. Right: merge of both channels with GFP-PH in green and mRuby-Lifeact in magenta. All movie timestamps are relative to anaphase onset (mm:ss).

Video S5: Simultaneous imaging of membrane and aPKC dynamics during neuroblast asymmetric division.

Time lapse imaging of Drosophila third instar larvae neuroblast expressing GFP-aPKC from its endogenous promoter and mCherry-PH via worniu-GAL4. Time relative (mm:ss) to nuclear envelope breakdown (NEB) is shown with a 20 seconds/frame time resolution. Left: GFP-aPKC. Middle: mCherry-PH. Right: merge of both channels with GFP-aPKC in green and mCherry-PH in magenta. Top row is maximum intensity projection (MIP) of the front hemisphere of the neuroblast. Bottom row is a medial section. As described previously, aPKC accumulates on the apical membrane beginning in early prophase followed by a coalescence phase shortly before NEB. Membrane features are present throughout the cell cycle, and their movements occur simultaneously with aPKC.

Video S6: aPKC is recruited to apical membrane patches

Time lapse imaging of Drosophila third instar larvae neuroblast expressing GFP-aPKC from its endogenous promoter and mCherry-PH via worniu-GAL4. Shown are maximum intensity projections (MIP) of the front hemisphere of the neuroblast. Left: GFP-aPKC. Middle: mCherry-PH. Right: merge of both channels with GFP-aPKC in green and mCherry-PH in magenta. Time relative (mm:ss) to nuclear envelope breakdown (NEB) is shown 20 seconds/frame time resolution. Depicted is an example of aPKC recruitment to a membrane patch as it moves from the basal to the apical hemisphere. Yellow arrowheads mark the membrane patch and the corresponding aPKC signal.

Video S7: Membrane and aPKC dynamics in a cyclodextrin-treated neuroblast

Time lapse imaging of membrane and aPKC dynamics in a Drosophila third instar larvae neuroblast in which membrane domains were removed via cyclodextrin treatment. Neuroblast expressed GFP-aPKC from its endogenous promoter and mCherry-PH via worniu-GAL4. Shown are maximum intensity projections (MIP) of the front hemisphere of the neuroblast. Left: GFP-aPKC. Middle: mCherry-PH. Right: merge of both channels with GFP-aPKC in green and mCherry-PH in magenta. Time relative (mm:ss) to nuclear envelope breakdown (NEB) is shown with a 20 seconds/frame time resolution.

Video S8: Membrane dynamics in a neuroblast expressing RNAi directed against aPKC

Time lapse imaging of a *Drosophila* third instar larval neuroblast expressing GFP-PH under conditions of RNAi mediated knockdown of aPKC via worniu-GAL4. Shown is a maximum intensity projection (MIP) of the front hemisphere of the neuroblast. 20 seconds/frame time resolution.

Video S9: Miranda dynamics in a cyclodextrin-treated neuroblast

Time lapse imaging a *Drosophila* third instar larval neuroblast expressing GFP-Miranda via worniu-GAL4 and RFP-H2A via a constitutive promoter. Two different imaging conditions were employed: untreated control (first movie) and cyclodextrin treatment (second movie). Shown is a maximum intensity projection (MIP) of the front hemisphere of the neuroblast. Left: GFP-Miranda. Middle: RFP-H2A. Right: merge of both channels with GFP-Miranda in green and RFP-H2A in magenta. Time relative (mm:ss) to anaphase onset is shown.

Video S10: Membrane and aPKC dynamics in a Latrunculin A treated neuroblast

Time lapse imaging a Drosophila third instar larvae neuroblast expressing GFP-aPKC via endogenous promoter and mCherry-PH via worniu-GAL4. Shown is a maximum intensity projection (MIP) of the front hemisphere of the neuroblast. Left: GFP-aPKC. Middle: mCherry-PH. Right: merge of both channels with GFP-aPKC in green and mCherry-PH in magenta. Movie A begins with the neuroblast being untreated before LatA was added to the media. Caption denotes when LatA was added. Time relative (mm:ss) to anaphase onset is shown 20 seconds/frame time resolution.

